# Classifying and Mapping Wetland Vegetation Assemblages in Coastal Louisiana with Landsat Imagery, 1985-2025

**DOI:** 10.64898/2026.08.13.744705

**Authors:** Gregg A. Snedden, Brady Couvillion, Donald R. Schoolmaster

## Abstract

The tidal wetlands of Louisiana comprise about 25% of those found throughout the conterminous United States yet estimates of wetland loss rates in the region between 1932 and 2016 have exceeded 60 km^2^ yr^-1^. To mitigate further degradation and wetland loss in the region, a globally unprecedented $50B, 50-year plan for coastal Louisiana is driving restoration efforts, and demand exists from multiple stakeholders for regularly updated, regional-scale, accurate land cover information. We used machine learning (random forests; RF) and cloud computing to develop a new Landsat-based, marsh vegetation community geospatial dataset. The dataset depicts wetland vegetation community types defined in a previous study at annual (1985–2025) time steps at 30-m resolution. An RF algorithm was used to integrate training samples with feature variables derived from Landsat imagery, and the resulting geospatial data product achieved an overall correct classification rate of 78%. The approach for development of the land cover dataset presented here has potential for application in other coastal wetland habitats throughout the world.

## Background & Summary

Coastal wetland ecosystems are expected to undergo significant hydrological changes under current environmental change projections, potentially leading to shifts in marsh vegetation species composition^1–4^. The processes driving these shifts include anticipated changes in salinity and inundation regimes driven by sea-level rise^5,6^ and changes in freshwater delivery that may arise from altered precipitation at regional and continental scales^7,8^.

Coastal Louisiana, situated in the south-central United States, contains approximately 40% of the coastal wetlands in the continental United States and has experienced some of the highest rates of land loss in North America, with over 5,000 km^2^ of coastal wetlands converting to open water over the last century^9^. This wetland loss has largely occurred in response to relative sea-level rise^10^ and reductions in sediment supply^11^ and, as such, the region is now the focus of a globally unprecedented ecosystem restoration program estimated to cost $50B over the next 50 years^12^. Wetland vegetation communities serve as sensitive indicators of ecosystem processes and environmental conditions across both local and landscape scales^13^, and various plant assemblages can influence key functions such as hydrodynamics^14^, vertical accretion^15^, and carbon burial^16^ in distinct ways. Consequently, long-term monitoring of marsh vegetation composition and distribution has been underway for over 30 years^17,18^, with continued observation used to assess ecosystem responses to restoration interventions and ongoing stressors such as sea-level rise and tropical cyclone landfalls^19^.

To date, vegetation monitoring in coastal Louisiana has principally included ground-based ocular species-cover surveys performed either periodically since 1993 to monitor specific restoration projects supported by the Coastal Wetlands Planning, Protection and Restoration Act (CWPPRA)^20^ or annually at 390 fixed monitoring sites as part of the Coastwide Reference Monitoring System (CRMS)^18^. CRMS is a systematic and extensive monitoring program established in 2006, which is supported by CWPPRA and by restoration programs created following the Deepwater Horizon Oil Spill of 2010, including the Louisiana Trustee Implementation Group for the Natural Resource Damage Assessment Trustees. In addition to the ground-based surveys, aerial surveys have been conducted late summer from a hovering helicopter at approximately 4,000 plots (30-m radius) situated along approximately 150 meridional transects in 1997, 2001, 2007, 2013, and 2021. Ordination and cluster analyses (two-way indicator species analysis [TWINSPAN])^21^ have been used to classify the 1997 and 2001 aerial survey data into distinct vegetation community assemblages^22–24^.

Though these efforts to monitor the distribution of emergent marsh vegetation taxa and classify sites into distinct assemblages provide information to evaluate and support restoration efforts, they also carry limitations and tradeoffs. The aerial surveys, while providing high spatial resolution, are irregularly spaced in time, with the time interval between surveys averaging six years. As such, they are unable to resolve interannual variations in vegetation distribution such as those that may arise from precipitation teleconnections with El Niño-Southern Oscillation (ENSO) activity^8^, which carries a recurrence interval of 2–7 years^25,26^. While the annual ground-based CRMS surveys address this limitation, their spatial resolution is far reduced (*n* = 390), as is the record length in that the CRMS monitoring program was not initiated until 2006. Until recently, classification of sites into distinct vegetation assemblages based on their species relative-cover vectors was principally conducted with parametric, distribution-based multivariate clustering approaches such as TWINSPAN. However, these approaches can exhibit poor stability when incorporating additional data such as those generated in long-term monitoring programs, as the cluster assignments of previously classified samples may change if the additional samples sufficiently alter the existing multivariate distribution of the data^27^. This lack of stability presents significant challenges to long-term monitoring programs, as it may undermine the ability to make meaningful temporal comparisons.

Recently, artificial neural network approaches have gained appeal as alternatives to distribution-based statistical methods^28^ for ecological assemblage delineation. Among them, self-organizing maps (SOMs)^29^ have been broadly applied to fish^30,31^, benthic macroinvertebrates^32,33^, forest vegetation^34^, and, recently, emergent marsh vegetation in coastal Louisiana^35^. After training an SOM classifier, new samples can be projected onto it for assignment to predefined assemblages or community types. This feature makes SOMs ideal for classifying data in long-term, ongoing ecological monitoring programs. The 11 emergent marsh vegetation communities defined in Snedden^35^ have since been incorporated into the CRMS program, and into modeling efforts that support decisions related to restoration planning in coastal Louisiana.

Remote sensing approaches offer significant advantages over *in-situ* approaches for mapping zonation of vegetation assemblages across extensive geographic areas, and have gained prominence due to their broad spatial coverage, temporal consistency, cost-effectiveness, and capacity for integration with other geospatial data for further analysis^35^. Common research themes, including mapping vegetation types and multitemporal change detection, have largely used medium spatial resolution data such as Landsat imagery due to its balance of temporal frequency (16-d), spatial resolution (30-m), and broad accessibility^37^. Numerous studies have demonstrated the utility of leveraging Landsat imagery for mapping wetland vegetation zonation with acceptable accuracy^38–41^. The evolution of remote sensing-based classification methods has encompassed a diverse array of algorithmic approaches, including supervised, unsupervised, and non-parametric techniques^42^. Among these techniques, supervised classification algorithms have been extensively implemented for land cover mapping applications in wetland settings^42–44^.

Despite the utility of these approaches, the inevitability of classification errors and inaccuracies associated with image classification necessitates rigorous accuracy assessment protocols for quantifying the reliability and statistical confidence of remotely sensed land-cover products.

The Louisiana Coastal Wetland Vegetation Community Dataset (LCWVCD) described here synthesizes (1) an existing SOM-based emergent marsh vegetation community classifier^35^, species-cover datasets from aerial surveys conducted in 1997^45^, 2001^46^, 2007^47^, 2013^48^, and 2021^49^, and (3) historical Landsat imagery (1985 to present) to classify and map emergent marsh vegetation community types at annual time steps and 30-m resolution. We first project all samples from the aerial surveys onto the SOM classifier to assign one of 11 distinct community types to each of the aerial survey samples. We then integrate the NOAA Coastal Change Analysis Program Regional Land Cover Dataset (C-CAP; https://coast.noaa.gov/digitalcoast/data/ccapregional.html) to add training and validation samples to classify mangrove forest regions. Next, half of the classified aerial survey data are used as targets for supervised classification of Landsat imagery to those 11 community types. Finally, after classifying the Landsat imagery, we use the validation data to assess accuracy of the classified Landsat imagery.

## Methods

### Study area

Coastal Louisiana, located between 29.0–30.5°N and 89.0–94.0°W, encompasses approximately 15,000 km^2^ of marshes, swamps, and lowland habitats that border the northern Gulf of America (Gulf of Mexico) forming a continuum of vegetation community types that largely reflect the region’s long-term estuarine salinity gradient (Fig. 1). The landscape is composed of two distinct geomorphic regions—the Mississippi River Deltaic Plain (MRDP) and the Louisiana Chenier Plain (LCP). The MRDP extends along eastern coastal Louisiana (east of 92°W) and sits atop Holocene sediment deposits that originated from the 3.2M km^2^ Mississippi River drainage basin and accumulated over 8,000 years of river avulsion events near the river’s mouth. West of the river’s mouth, estuary-ocean exchange occurs primarily through a series of tidal passes situated between barrier islands. The LCP, located in southwestern Louisiana (west of 92°W) was also formed with alluvial materials originating from the Mississippi River drainage basin, but after they were discharged to the coastal ocean, carried westward by a nearshore coastal current, and subsequently reworked against the coast through wave action^50,51^. The LCP features ridges (cheniers) and levees that tend to restrict tidal exchange with the coastal ocean to a few narrow inlets that incise the ridges. The reduced tidal exchange in this region is highly consequential to the region’s hydrology in that it can lead to hypersaline conditions during drought^8^.

**Fig 1.**
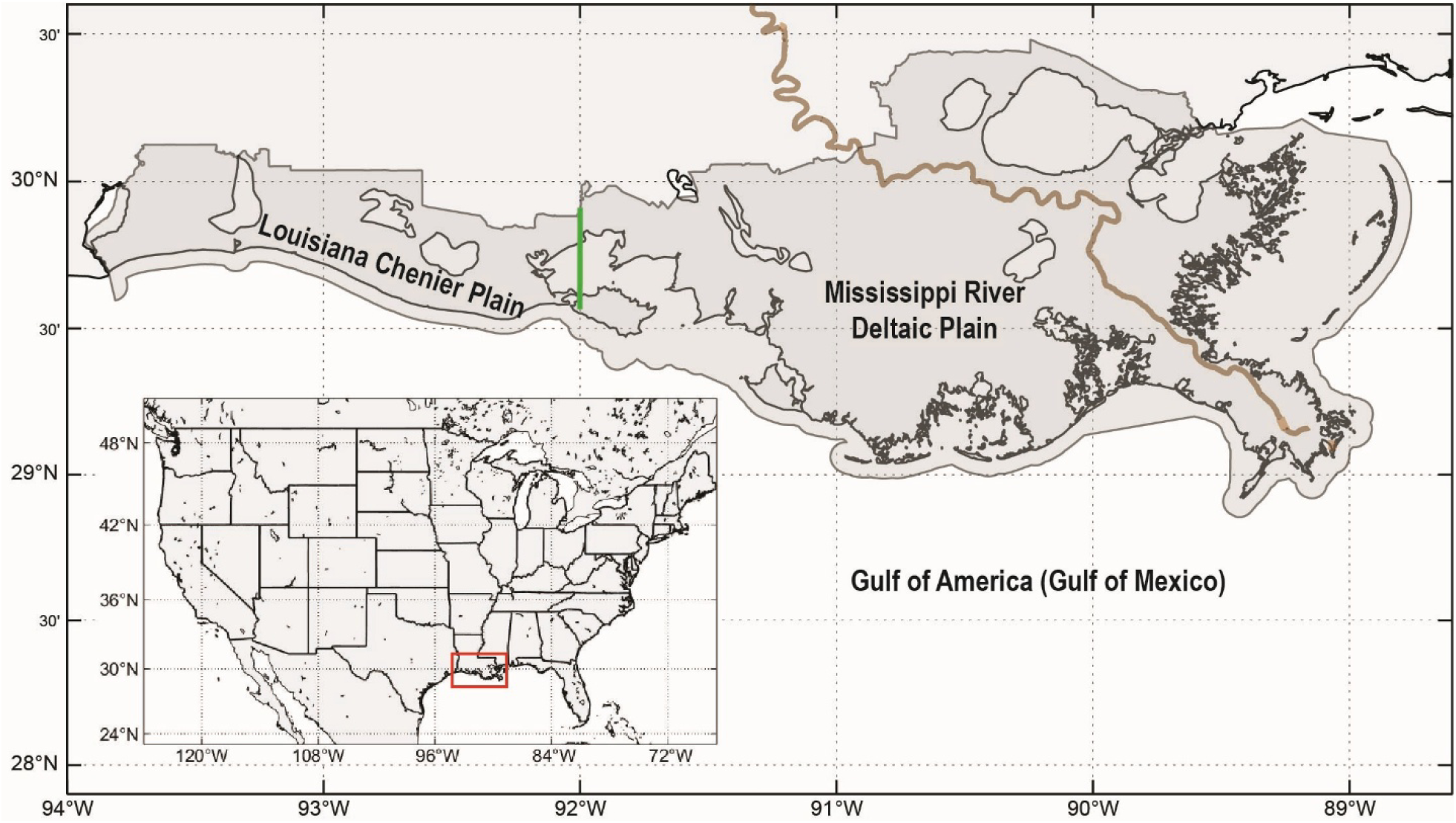
Map of the study area. The coastal zone, the region over which the land-cover dataset was produced, is shaded gray. The Mississippi River is shown by the brown line. The green vertical line separates the Louisiana Chenier Plain (west) from the Mississippi River Deltaic Plain (east). The inset shows the location of the study area (red box) within the conterminous United States.

### Overview of Workflow

The workflow for generation of the Louisiana Coastal Wetland Vegetation Community Dataset (LCWVCD) included (1) SOM training to define and delineate vegetation community types, (2) training/validation sample generation, (3) image processing, (4) feature variable generation, (5) model training with random forests (RF), (6) application of RF model to feature variable composites for generation of annual land cover products, and (7) validation and accuracy assessment (Fig. 2). Image processing, feature variable generation, and RF training were primarily conducted on the Google Earth Engine cloud-based geospatial platform.

**Fig 2.**
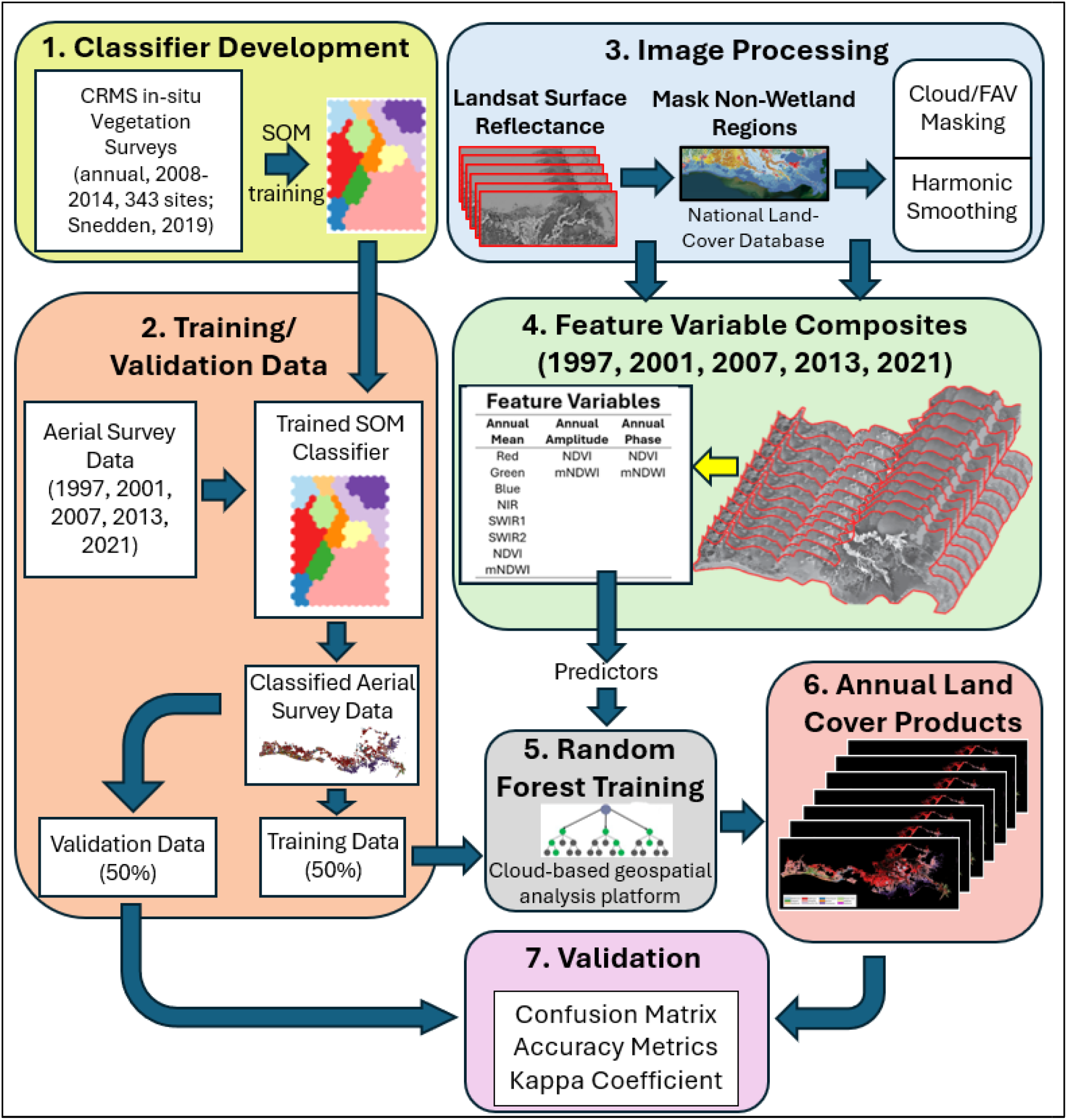
Overall workflow for generation of the Louisiana Coastal Wetland Vegetation Community Dataset (LCWVCD).

### Delineation of vegetation assemblages/classifier development

A SOM was trained with data from 2,526 *in-situ* emergent marsh vegetation surveys that assessed relative cover of 49 taxa that were performed annually 2008–2014 at 343 CRMS sites (each consisting of ten 2-m × 2-m survey plots randomly located along a 200-m transect). To train an SOM, a multidimensional dataset—such as a species relative cover dataset assembled from surveys conducted at multiple sites and/or multiple sites—is fed into a nonlinear ordination algorithm and projected onto a two-dimensional ordination space. The ordination space consists of a collection of discrete cells, and each cell has a species composition. Like linear approaches to ordination, samples projected onto SOM cells near each other in the ordination space have similar species compositions; those projected far apart have different species compositions. A SOM cell-by-species matrix is assembled, to which cluster analysis is performed to assign each cell a cluster membership (Fig. 2, box 1). After ordination and clustering, new samples (not used for SOM training) can be projected onto the SOM by identifying the cell whose species composition minimizes the Euclidean distance (the best-matching unit; BMU) to that of the new sample being classified, and cluster membership is assigned to it such that it reflects the BMU’s cluster membership. The resulting SOM classifier was a 260-cell lattice consisting of 20 rows and 13 columns. Eleven marsh vegetation community types were delineated (maidencane; bulltongue; three-square; roseau cane; paspalum; wiregrass; bulrush; saltgrass; needlerush; brackish mix; oystergrass).

Dominant taxa and mean relative % cover values for each community type (2014–2025) are depicted in Fig. 3 (left), along with swarm plots of their mean salinity and % time inundated (October, 2021–September, 2022) based on each vegetation plot’s water-surface elevation and salinity time series data, and each vegetation plot’s elevation data (https://www.lacoast.gov/crms_viewer/Map/CRMSViewer). Further details related to SOM training and descriptions of the resulting community types are provided in Snedden^35^.

**Fig 3.**
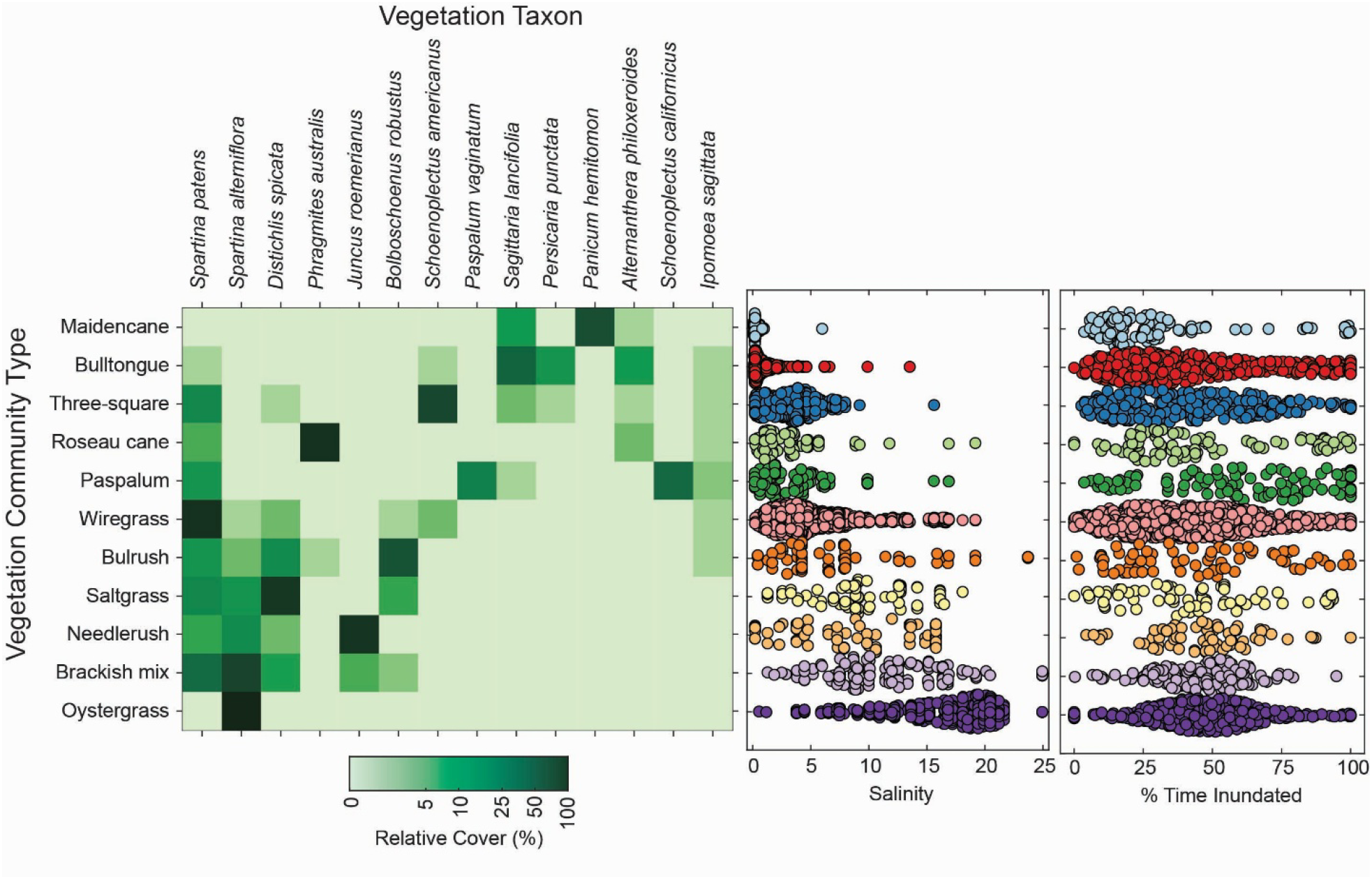
(left) For each vegetation community type, the mean relative percent cover of the 14 vegetation taxa most important to structuring the self-organizing map classifier, for all Coastwide Reference Monitoring System (CRMS) *in-situ* vegetation plots sampled during 2014–2025 water years. (center/right) Distribution of average salinity (center) and percent time inundated (right) during the October, 2021–September, 2022 water year for all CRMS *in-situ* vegetation plots in each vegetation community type.

### Training/validation data

Helicopter-based marsh vegetation survey data from 1997^45^, 2001^46^, 2007^47^, 2013^48^, and 2021^49^—each consisting of approximately 4,000 plot surveys—were truncated to the 49 taxa included in the SOM classifier, after which relative cover values were calculated for each sample. Each helicopter-based relative cover sample was then projected onto the SOM to assign to it one of the 11 marsh vegetation community types (Fig. 2, box 2). In addition to sites with marsh vegetation cover, during each survey year approximately 100 training/validation points were placed in regions classified by the C-CAP Regional Land Cover Dataset (NOAA) as “estuarine shrub-scrub” and assigned as ‘mangrove’. For each survey year, 50% of the training/validation data were randomly selected from each vegetation community type for RF training, and the remaining 50% were set aside for validation after Landsat imagery was classified.

### Landsat imagery processing and feature variable generation

Landsat Collection 2 surface reflectance data^52–54^, which provide spatially continuous coverage of the earth’s surface from 1985 to 2025 at 30-m spatial resolution with a 16-d flyover return period, were acquired from the U.S. Geological Survey Earth Explorer (https://earthexplorer.usgs.gov/). Time series for various spectral bands, including blue (450–510 nm), green (530–590 nm), red (640–670 nm), near infrared (NIR; 850–880 nm), shortwave infrared 1 (SWIR1; 1570–1650 nm), and shortwave infrared 2 (SWIR2; 2110–2290 nm) were compiled, after which clouds and cloud shadows were flagged for exclusion via codes embedded in the pixel quality band. For each band, cloud-contaminated pixels were imputed with 35-d moving median values (Fig. 2, box 3).

The modified normalized difference water index (mNDWI)^55^, which enhances water features while reducing noise from land, vegetation and soil, and was calculated for each cloud-free pixel as

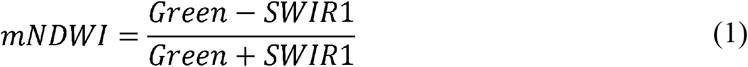

and used to facilitate the differentiation between true land and floating aquatic vegetation (FAV; details provided below). Normalized difference vegetation index (NDVI)^56,57^ was calculated for each cloud-free pixel as

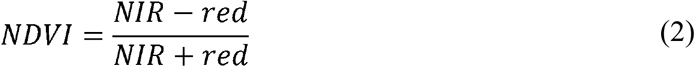

and used as a proxy for the presence and vigor of vegetation.

After cloud masking, temporal imputation of cloud-contaminated pixels, and calculation of mNDWI and NDVI for each pixel, each pixel’s annual (Sep–Aug) time series for reflectance in each of the six spectral bands and the two derived indices (NDVI; mNDWI) were fit with annual harmonics for temporal smoothing. Annual means for each of the eight smoothed time series were calculated and retained, along with the amplitude and phase of Fourier-smoothed NDVI and mNDWI time series. In this fashion, each pixel carried 12 feature variables for each year, which were subsequently used as independent variables in RF training (Fig. 2, box 4).

Finally, for each year, each pixel’s red and NIR spectral reflectance bands and two derived indices (NDVI and mNDWI) were aggregated into (separate) seasonal (winter [DJF], spring [MAM], summer [JJA] and fall [SON]) composites and, for each season in each year, medians and standard deviations were derived for each of the four metrics. The variation in the indices among the seasonal composites were used to mask out FAV; (refer to ref [58] for detailed methods). The resulting land/water/FAV dataset was further composited with categories from the annual National Land Cover Database (NLCD)^59^ to further assign pixels to land-cover classes such as uplands, emergent herbaceous wetlands, developed areas, bottomland hardwood forests, and others. Only pixels classified as emergent herbaceous wetlands in the resulting dataset were retained and classified further into the 11 marsh vegetation/mangrove community types.

### Model training and prediction with random forests

This study utilizes RF algorithms in Google Earth Engine for model training. RF classifiers have gained widespread adoption for land cover classification applications due to their capacity to meet challenges of high-dimensionality and nonlinearity while minimizing overfitting issues^60,61^. In land-cover mapping exercises, RF algorithms exhibit superior performance compared to other machine learning approaches and, additionally, they exhibit resilience to image noise, making them well-suited for remote sensing applications. Prior studies have shown that using a large number of trees (>50) produces increased accuracy for land-cover classification applications^62^, with diminishing returns once the number of decision trees surpasses 100. Consequently, 100 trees were selected for this investigation. Additionally, we set the minimum leaf population to two, which allows for precise distinctions between land-cover classes while mitigating the risk of overfitting. We used default values for all remaining settings. For each of the five helicopter survey years, the 12 variables in the annual feature variable composites were used to create a predictor dataset composed of 12 feature bands to train an RF classifier (Fig. 2, box 5), after which model parameters for each of the five resulting RF classifiers were averaged to create a composite RF classifier. This composite RF classifier was then applied to each of the 1985–2024 annual feature composites to produce land-cover data products (Fig. 2, box 6).

#### Data Records

The LCWVCD provides annually updated geospatial layers of vegetation community types from 1985 to 2025 at 30-m spatial resolution and publicly available in GeoTIFF format^63^. Each pixel carries an integer value that corresponds to one of 26 land cover classes (Table 1). Pixel values 13–24 indicate one of the 11 herbaceous marsh vegetation/mangrove classes described above. Pixel values 1–12, and 25 correspond to non-wetland vegetation classes. Pixel value 26 corresponds to open water.

**Table 1.** List of pixel values assigned to each land cover class. Pixel values 13–24 indicate one of the wetland vegetation community types described above and in Snedden^35^; pixel values 1–12 indicate land cover classes other than herbaceous wetland/mangrove communities; pixel values 25 and 26 indicate floating aquatic vegetation and open water, respectively.

| Value | Land Cover Class | Value | Land Cover Class |
| --- | --- | --- | --- |
| 1 | Developed, High Intensity | 14 | Bulltongue |
| 2 | Developed, Medium Intensity | 15 | Three-Square |
| 3 | Developed, Low Intensity | 16 | Roseau Cane |
| 4 | Developed, Open Space | 17 | Paspalum |
| 5 | Sand | 18 | Wiregrass |
| 6 | Barren Land | 19 | Bulrush |
| 7 | Upland Forest | 20 | Saltgrass |
| 8 | Shrub/Scrub | 21 | Needlerush |
| 9 | Grassland/Herbaceous | 22 | Brackish Mix |
| 10 | Pasture/Hay | 23 | Oystergrass |
| 11 | Cultivated Crops | 24 | Mangrove |
| 12 | Woody Wetlands | 25 | Floating Aquatic Vegetation |
| 13 | Maidencane | 26 | Open Water |

#### Data Overview

Over 75% of total wetland cover in the Louisiana coastal zone from 1985–2025 consisted of either maidencane, bulltongue, wiregrass, or oystergrass communities (Table 2). Bulltongue communities exhibited no significant change in coverage over time, while coverage of maidencane and wiregrass communities has been significantly declining. Pixels classified as non-permeant water (i.e., those that were not ‘open water’ for the entire 41-year record such as the coastal ocean, lakes, or streams) have been increasing at a rate of 29.1 km^2^ yr^-1^, which corresponds well to the 1984–2020 land loss estimate of 34.8 km^2^ yr^-1^ put forth in Jenson et al.^64^. The Delta Plain in southeastern Louisiana, which exhibits tidal connectivity with the coastal ocean, shows a general down-estuary progression from maidencane to bulltongue to wiregrass to brackish mix to oystergrass (Fig. 4), largely reflecting the estuarine salinity gradient (Fig. 3, center panel). Brackish mix and oystergrass communities are largely absent from the Chenier Plain in southwestern Louisiana, possibly a reflection of the reduced tidal exchange in that region.

**Table 2.** Area, expressed in km^2^ and percentage of marsh/non-permanent water cover classes, and time rate of change, determined by linear regression, for the various marsh vegetation community types. Non-permanent water category includes pixels that were classified as one of the marsh vegetation community types for some years, and water for other years, 1985–2025.

| Wetland Cover Class | mean area ($\text{km}^2$) | % of total wetland area | slope ($\text{km}^2 \text{ yr}^{-1}$) | slope SE ($\text{km}^2 \text{ yr}^{-1}$) | $r^2$ | $p$ |
| --- | --- | --- | --- | --- | --- | --- |
| Maidencane | 898 | 6.8 | -10.5 | 1.9 | 0.43 | <0.001 |
| Bulltongue | 2878 | 21.7 | -4.7 | 3.4 | 0.02 | 0.173 |
| Three-Square | 648 | 4.9 | +19.2 | 2.0 | 0.69 | <0.001 |
| Roseau | 368 | 2.8 | +3.0 | 0.8 | 0.25 | 0.001 |
| Paspalum | 332 | 2.5 | +7.3 | 1.1 | 0.51 | <0.001 |
| Wiregrass | 4373 | 32.9 | -52.8 | 4.0 | 0.81 | <0.001 |
| Bulrush | 40 | 0.3 | +2.2 | 0.3 | 0.60 | <0.001 |
| Saltgrass | 67 | 0.5 | +1.1 | 0.5 | 0.11 | 0.022 |
| Needlerush | 199 | 1.5 | +3.1 | 0.6 | 0.39 | <0.001 |
| Brackish Mix | 677 | 5.1 | +1.7 | 1.9 | 0.00 | 0.376 |
| Oystergrass | 701 | 5.3 | +1.9 | 2.1 | 0.00 | 0.367 |
| Mangrove | 40 | 0.3 | -0.6 | 0.3 | 0.06 | 0.069 |
| Non-Permanent Water | 2067 | 15.6 | +29.1 | 1.8 | 0.86 | <0.001 |

**Fig 4.**
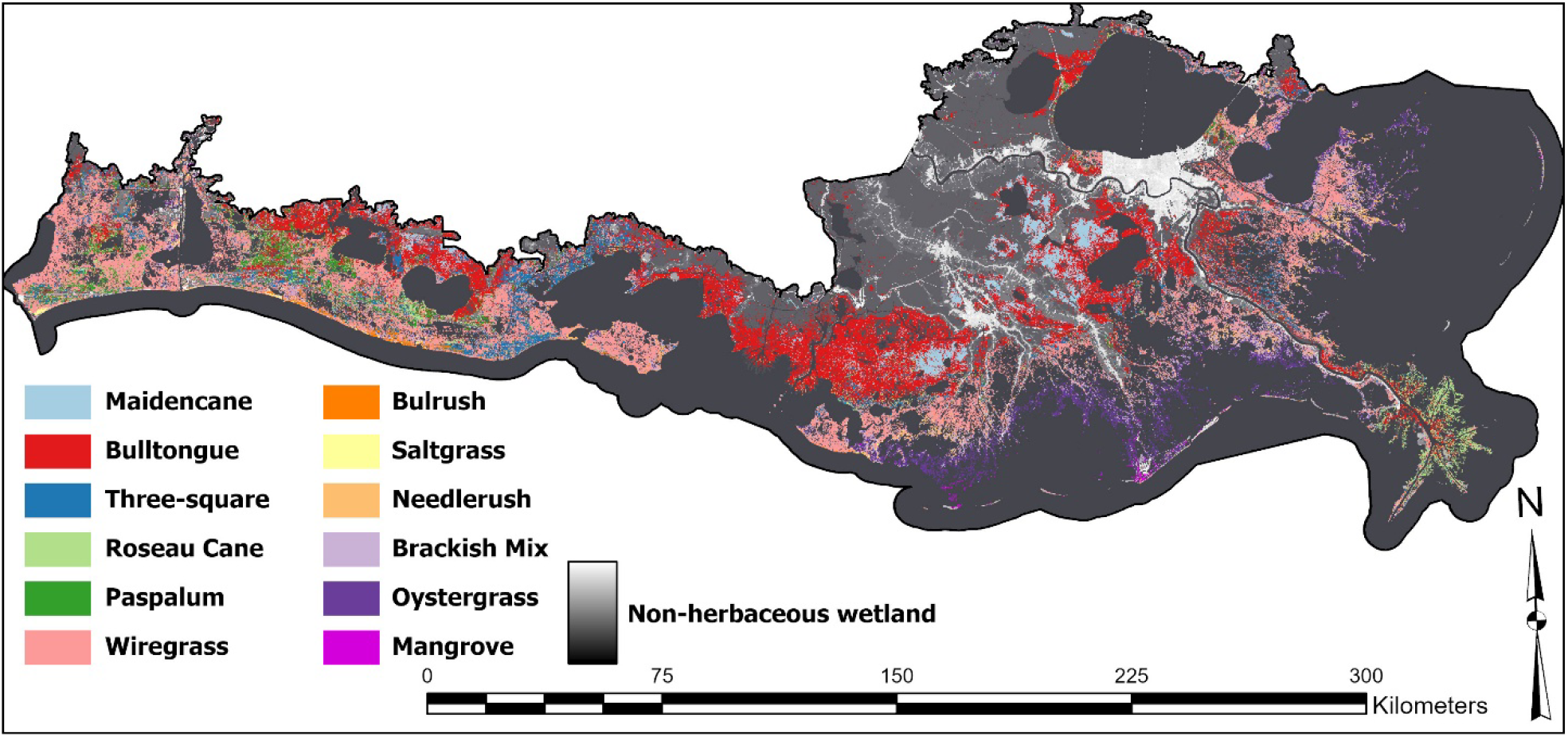
Vegetation community land cover map of the Louisiana coastal zone in 2024.

#### Technical Validation

The remaining half of the classified helicopter-based surveys (*n* = 9,890) were used with their corresponding (in time and space) Landsat pixel classifications to assemble a confusion matrix for each year. The five confusion matrices were aggregated into one (Table 3), which was used as the basis for accuracy validation. The validation metrics included producer’s accuracy (PA; Table 4), user’s accuracy (UA), overall correct classification rate (CCR), and the kappa coefficient^65^.

**Table 3.**
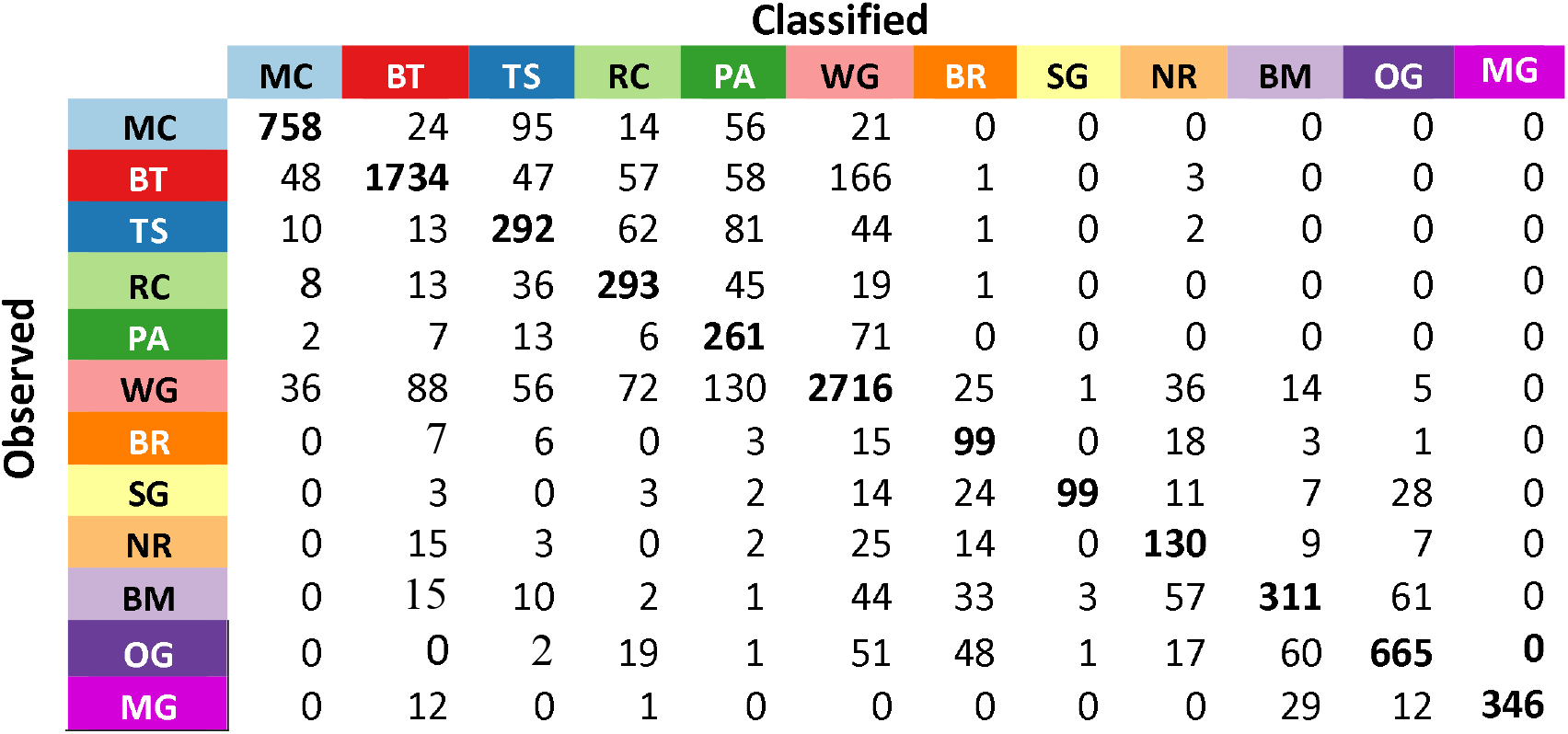
Confusion matrix for validation data. Abbreviations are MC (maidencane); BT (bulltongue); TS (three-square); RC (roseau cane); PA (paspalum); WG (wiregrass); BR (bulrush); SG (saltgrass); NR (needlerush); BM (brackish mix); OG (oystergrass); MG (mangrove). Bold numbers along matrix diagonal are counts of correct classifications. Off-diagonal counts indicate misclassifications.

**Table 4.** Accuracy metrics for the land cover classes. Numbers below community types indicate percentage of validation samples that fall under each land-cover classification. Abbreviations are MC (maidencane); BT (bulltongue); TS (three-square); RC (roseau cane); PA (paspalum); WG (wiregrass); BR (bulrush); SG (saltgrass); NR (needlerush); BM (brackish mix); OG (oystergrass); MG (mangrove).

| Accuracy Metric | MC (10) | BT (21) | TS (5) | RC (4) | PA (4) | WG (31) | BR (2) | SG (2) | NR (2) | BM (5) | OG (9) | MG (5) |
| --- | --- | --- | --- | --- | --- | --- | --- | --- | --- | --- | --- | --- |
| Producer's | 0.78 | 0.82 | 0.58 | 0.71 | 0.73 | 0.85 | 0.65 | 0.52 | 0.63 | 0.58 | 0.77 | 0.87 |
| User's | 0.88 | 0.90 | 0.52 | 0.55 | 0.41 | 0.85 | 0.40 | 0.95 | 0.47 | 0.72 | 0.85 | 1.00 |

The CCR of the resulting aggregated (1997, 2001, 2007, 2013, 2021) confusion matrix, which reflects the proportion of correctly classified pixels across all community types, was 78%, with a kappa coefficient of 0.73, indicating ‘substantial agreement’^66^. The confusion matrix shows elevated occurrences of misclassifications of (1) brackish mix into wiregrass, bulrush, needlerush, and oystergrass communities; (2) three-square communities into Roseau cane, paspalum, and wiregrass communities; and (3) saltgrass communities into bulrush and oystergrass communities. Each of these groups of communities shared common sets of dominant taxa, but with differing relative dominance among the taxa (Fig.3, left panel). While clustering the SOM’s cell-by-species matrix produces crisp, distinct boundaries around the assemblages, transitions in relative cover among the vegetation taxa through space and time are fuzzy and gradual^67^. The geographic zonation of the community types generally aligns with the estuarine salinity gradient in coastal Louisiana^35^ and as such, they are ordered in the confusion matrix by increasing salinity. Thus, the preponderance of zeros (or relatively small numbers) in the upper right and lower left regions of the confusion matrix indicates that misclassifications do not typically result in class assignments to community types far along the salinity gradient from the community type to which they should be correctly assigned.

## Data Availability

The dataset described in this study is publicly available at https://doi.org/10.5066/P13AKBYE

## Code Availability

The GEE scripts used to process Landsat imagery and train the RF classifier are available to the public at https://doi.org/10.5066/P13AKBYE.

